# Directional Absolute Coherence: a phase-based measure of effective connectivity for neurophysiology data

**DOI:** 10.1101/2022.02.07.479359

**Authors:** Maximilian Scherer, Tianlu Wang, Robert Guggenberger, Luka Milosevic, Alireza Gharabaghi

## Abstract

Communication between neural structures is a topic of much clinical and scientific interest and has been linked to a variety of behavioural, cognitive, and psychiatric measures. Here, we introduce a novel effective connectivity measure, termed the *directional absolute coherence* (DAC). Combining aspects of magnitude squared coherence, imaginary coherence, and phase slope index, DAC provides an estimate of connectivity that is resistant to volume conduction, encapsulates the directionality of neural communication, and is bound to the interval of –1 and 1. To highlight the properties of this newly proposed method, we compare DAC to a number of established connectivity methods using data recorded from the subthalamic nucleus of patients with Parkinson’s disease with deep brain stimulation electrodes. By applying a combination of real and simulated data, we demonstrate that DAC provides a reliable estimate of the magnitude and direction of connectivity, independent of the phase difference between brain signals. As such, DAC facilitates a reliable investigation of inter-regional neural communication, rendering it a valuable tool for gaining a deeper understanding of the functional architecture of the brain and its relationship to behaviour and cognition. A Python implementation of DAC is freely available at https://github.com/neurophysiological-analysis/FiNN.

**Highlights:**

- Volume conduction and limited interpretability affect many connectivity methods.
- DAC is a combined approach to overcome these pitfalls.
- DAC augments information content and interpretability in comparison to other methods.
- DAC allows for reliable estimation of effective connectivity.

## Introduction

The characterization of neural communication across different spatial and temporal scales opened up a whole new area of neuroscientific research (e.g., Bressler, 1995; Bullmore and Sporns, 2009). It has played a fundamental role in linking neural connectivity to a variety of behavioural and cognitive measures (e.g., Gaudet et al., 2020; Imperatori et al., 2014; Uhlhaas and Singer, 2006; Qi et al., 2020).

In the motor system, for example, long distance neural communication is essential in both the healthy and the pathological condition. In the healthy condition, active and passive movements show different patterns of communication between the brain and the periphery, as revealed by changes of cortico-muscular and muscular-cortical connectivity (Bourguignon et al., 2014; for a review, see e.g., Lattari et al., 2010). Pathological conditions, such as Parkinson’s disease (PD), also alter long-distance neural communication. In PD patients, the connectivity between the primary motor cortex and the subthalamic nucleus (STN) is increased (Baudrexel et al., 2011), and varies during motor execution (Lalo et al., 2008, Litvak et al., 2012), movement planning (Alhourani et al., 2020), as well as during rest (Kern et al., 2016; Belardinelli et al., 2019). Moreover, the intensity of information flow between the STN and the muscle differs between tremor-dominant from non-tremor PD patients, and correlates with tremor activity during voluntary movement (Naros et al., 2018). For this kind of neuroscientific insight with clinical relevance, the interaction between neural sources (e.g., cortical, subcortical and muscular) needs to be accurately captured.

In this context, one way of quantifying the communication between neural structures is to estimate the degree to which same-frequency neural oscillations relate to each other. This type of communication, referred to as same-frequency coupling, suggests that two neural structures communicate with each other, when oscillations in the same frequency interact with a consistent, non-zero phase shift (Fries, 2005). Same-frequency coupling is commonly described by the terms *functional* or *effective* connectivity (Friston, 2011). Functional connectivity refers to the temporal correlation between two spatially remote neurophysiological measurements (Friston et al., 1993). Effective connectivity expands upon this concept by including an estimate of the sequence in which oscillations are observed in different regions (Friston, 2011).

Given the broad fundamental and clinical interest in same-frequency coupling, many methods have been developed in a bid to accurately identify and quantify existing neuronal interactions while avoiding spurious connectivity patterns due to, e.g., noise or input from a common source (for reviews, see e.g., Bastos and Schoffelen, 2016; Bakhshayesh et al., 2019; Sakkalis, 2011). A large number of these methods rely on the coherency coefficient, which is a frequency domain equivalent of the time domain correlation coefficient (Nunez et al., 1997). Examples include magnitude squared coherence (Srinivasan et al., 1998) and phase lag index (PLI; Stam et al., 2007). These methods have been successful in revealing relevant brain-behaviour relationships (e.g., Litvak et al., 2012), however, numerous limiting factors and potential pitfalls have been highlighted and discussed in previous studies (e.g., Bakhshayesh et al., 2019; Li et al., 2020). One main challenge in estimating connectivity is the fact that activity from a single neural source can be observed in multiple sensors due to the spatial spread of electrical fields. This common phenomenon is called *volume conduction* (Nunez et al., 1997; Srinivasan et al., 2007), and may lead to erroneous estimates of connectivity between neighbouring sensors. Methods designed to overcome these artefacts include imaginary coherence (Nolte et al., 2004), phase slope index (PSI; Nolte et al., 2008), and weighted phase lag index (wPLI; Vinck et al., 2011). Of these methods, only the PSI is a measure of effective connectivity. However, one drawback of the PSI is that the outcome value is unbound. Nolte and colleagues therefore recommended that the PSI be normalized to its standard deviation (Nolte et al., 2008). Accordingly, multiple segments of PSI estimates need to be calculated if normalization is to be applied, necessitating a much larger volume of data than required with the other connectivity methods. This would limit its applicability in various situations where data collection is difficult, e.g., when acquiring data in patient populations in intraoperative settings.

This work introduces a novel measure of effective connectivity termed the *directional absolute coherence* (DAC). DAC combines the advantages of magnitude squared coherence, imaginary coherence, and PSI to overcome the disadvantages of each individual connectivity method. The resulting method presents a coherence-based measure that is directionalized, bound to the absolute interval of –1 to +1, robust towards volume conduction, and easily interpretable. In the next section, we begin with a description of the proposed method, and then illustrate its functionality in comparison to established connectivity methods using a mixed data set of real and simulated local field potentials.

## Methods

### Directional Absolute Coherence (DAC)

Conceptually, DAC consists of three distinct components which are combined to obtain a reliable estimate of the effective connectivity between two signals.

The first component is designed to determine the magnitude of connectivity strength between two signals. This may be done via magnitude squared coherence (Srinivasan et al., 1998), which is a functional connectivity measure which can be interpreted as the Pearson’s correlation between the spectral components of two signals and which is therefore bound to the interval of 0 and 1 (Equation 1A).

The second component is designed to detect the presence of volume conduction between the two signals. We based this component on imaginary coherence (Equation 1B; Nolte et al., 2004). Generally, imaginary coherence is calculated for single frequency bins and returns a value close to 0 when the phase shift between two signals approaches 0°; although zero crossings in imaginary coherence occur every 180°, these happen at different temporal lags for different frequencies. Hence, only for true volume conduction, the imaginary coherence across a range of frequencies is zero, whereas for false positives, it is only zero for a tiny minority. We utilize this property by counting the number frequency bins which are close to zero, below a specific threshold (e.g. 10°). Conceptually, selecting a threshold closer to 0° increases the sensitivity of the method towards connectivity at the expense of a decreased sensitivity towards volume conduction. If more than a specific percentage (e.g. 30%) of the values exceed the threshold, it indicates that the phase shift between the two signals is consistently close to zero across all evaluated frequency bins. In that case, the level of connectivity cannot be determined as volume conduction is certain. In our implementation of DAC, the function returns NaN (Not a Number – a common error value) when the phase shift between the two signals is less than 10° in more than 30% of the frequency bins. The threshold can be set arbitrarily and depends on the frequency investigated, distance between two sensors, and desired level of conservatism in detecting volume conduction. In the supplementary document, we provide an evaluation suite which can be used to configure said parameters. This evaluation suite can be executed in conjunction with a recording of one’s investigation in order to determine the ideal parameter configuration set.

The third component is designed to determine the directionality of the interaction between two signals. We selected the PSI (Equation 1C; Nolte et al., 2008), as it provides a robust measure of the direction of information flow from electrophysiological data with low signal-to-noise ratios (Haufe et al., 2011). Since we do not need a bounded value of PSI at this stage, no normalization is required. To encapsulate the directionality, DAC is multiplied by the sign of the unbounded output of PSI.

In summary, DAC is constructed by multiplying the magnitude squared coherence with an imaginary coherence-based estimate of volume conduction presenceand the sign of the PSI (Equation 1D). As a result, DAC is a measure of effective connectivity that is simple to interpret, statically bound within the interval –1 and +1, and robust towards volume conduction. A Python implementation of DAC can be found at https://github.com/neurophysiological-analysis/FiNN.

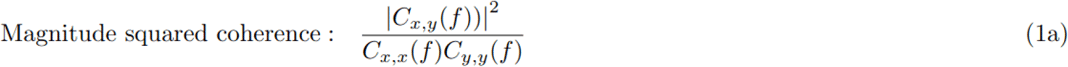

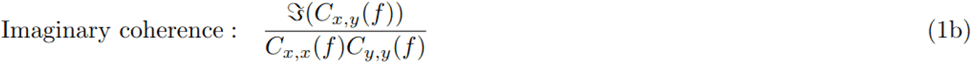

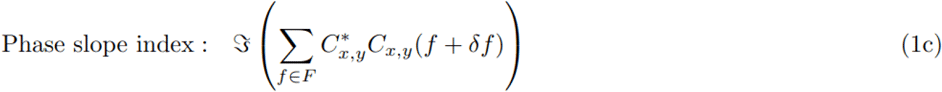

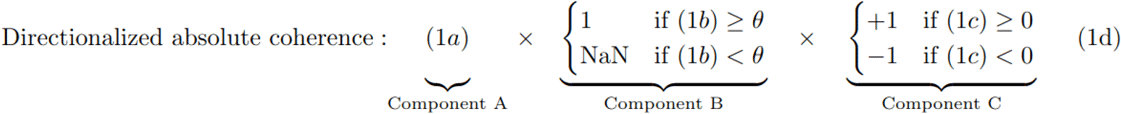

**Equation 1**. Measures of connectivity between input signals x and y. (**A**) Calculation of magnitude squared coherence. C_x,y_(f) denotes the complex cross-spectral density between signals x and y at frequency f. (**B**) Calculation of imaginary coherence. 𝕴(z) denotes the imaginary part of z. (**C**) Calculation of phase slope index. Δ denotes the frequency offset of the subsequent frequency bin. C*_x,y_(f) denotes the complex conjugate of the cross-spectral density between signals x and y at frequency f. θ is the threshold value for the phase shift angle between the two signals, below which volume conduction is detected. (**D**) Calculation of directed absolute coherence. Component A estimates the strength of the spectral relationship between two signals. Component B estimates the presence of volume conduction. Component C estimates the direction of the causal influence between two signals.

### Validation data

DAC was evaluated in comparison to magnitude squared coherence (Srinivasan et al., 1998), imaginary coherence (Nolte et al., 2004), normalized PSI (Nolte et al., 2008), and wPLI (Vinck et al., 2011). The reader may refer to the corresponding articles for a detailed description of the respective connectivity measures. To evaluate DAC in comparison to the aforementioned connectivity measures, we used exemplary data from a larger, previously published data set described in detail elsewhere (Milosevic et al., 2020). Briefly, four monopolar local field potential recordings were acquired from three patients with Parkinson’s disease during surgery for deep brain stimulation (Figure 1). We selected one recording from both Patient 1 and Patient 2 (recordings 1 and 2), and two recordings from two neighbouring lead contacts from Patient 3 (recordings 3a and 3b). The four 30-second recordings of each data set were selected on the basis of the magnitude of the peak in the beta frequency band in the power spectrum density.

**Figure 1.**
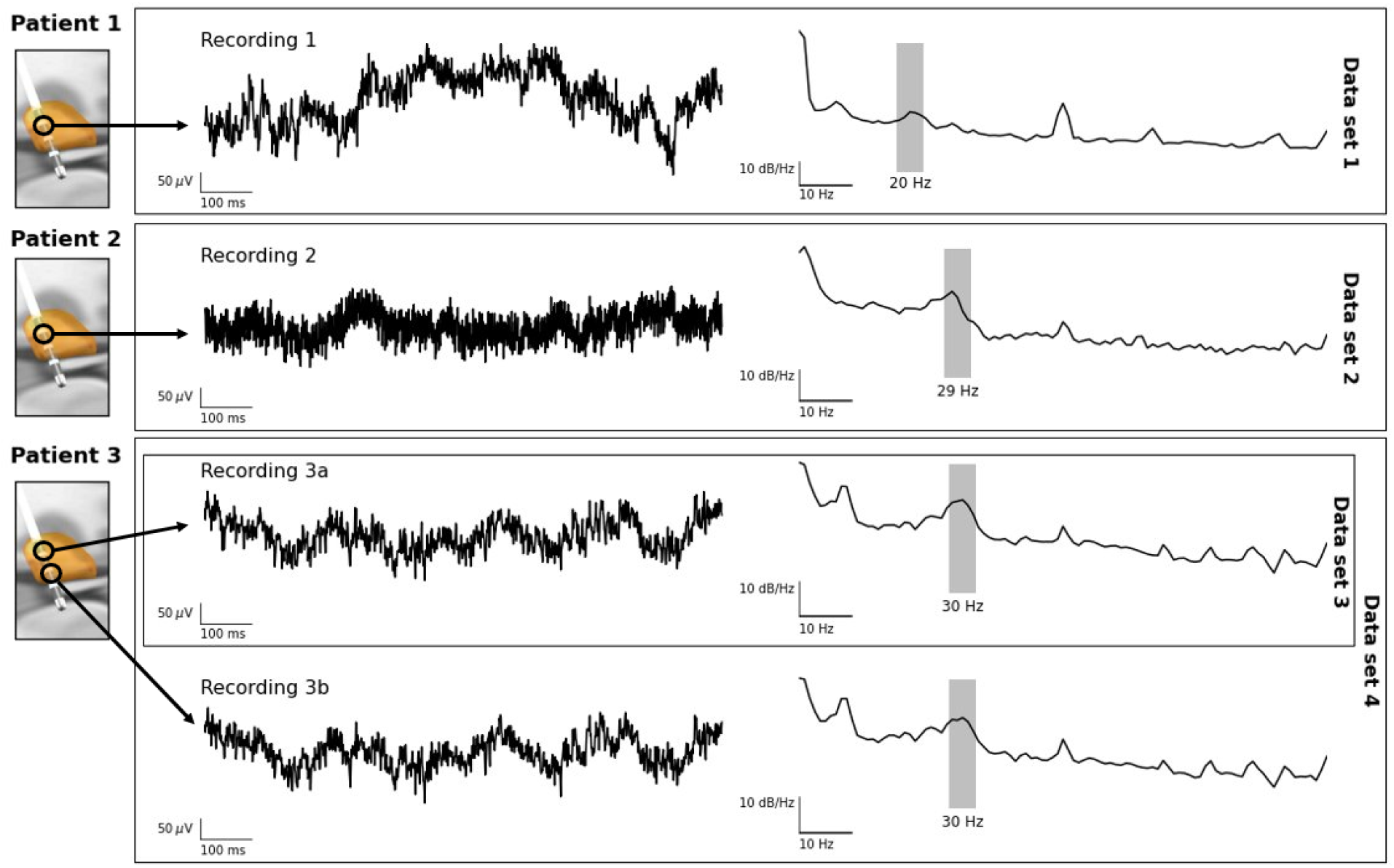
Illustration of human data used in validation. *Left*: Time domain representations of the signals used for the validation of the directed absolute coherence. For visualization purposes, only one second of data is shown for each recording. Recordings 1 and 2 were extracted from Patients 1 and 2, and used for data sets 1 and 2. Recordings 3a and 3b were extracted from two neighbouring deep brain stimulation contacts in Patient 3. Recording 3a was used for data set 3, while both recordings 3a and 3b were used for data set 4. *Right*: Power spectra of the four recordings. The areas shaded in grey mark the peak frequencies in the beta band; these are typical for data recorded from the subthalamic nucleus of patients with Parkinson’s disease.

To maintain full control of the level of expected connectivity, we used these recordings to construct four datasets that were based on real and simulated data. Specifically, recordings 1, 2 and 3a were used to construct data sets 1, 2 and 3, and the connectivity was calculated between the original signal and time-lagged versions of the same signal. In these three data sets, the signals therefore had identical background noise.

To construct data set 4, we used the recordings 3a and 3b (i.e., a pair of recordings from two neighbouring lead contacts), and the connectivity was calculated between the original signal 3a and the time-lagged versions of signal 3b. This approach enabled us to investigate signals that contained slightly different background noise.

The amount of time-lag was determined on the basis of the cycle length of the peak frequency in the beta frequency band. For example, in recording 1 with a peak at 20 Hz (Figure 1), the cycle duration was 50 ms. A wide range of time-lags was applied to the signal, corresponding to phase shifts from –270° to +270° in relation to the full cycle duration, in steps of 2°. For recording 1, these time-lags ranged from - 37.5 ms to +37.5 ms, in steps of 0.14 ms (= 50 ms / 360°). Using this approach, the time-lagged signal would precede the original signal for negative lags, simulating a flow of information from the time-lagged signal to the original signal. For positive lags, the time-lagged signal would follow the original signal, and the information flow would be reversed. The difference in time-lag may be interpreted as the difference in distance between the two signals. When the time-lag was sufficiently close to a duration corresponding to a 0° phase shift, we assumed a volume conduction effect between the two signals.

The scripts for this analysis can be found at https://github.com/VoodooCode14/dac.

## Results

Figure 2 shows the estimated connectivity values for different phase shifts.

**Figure 2.**
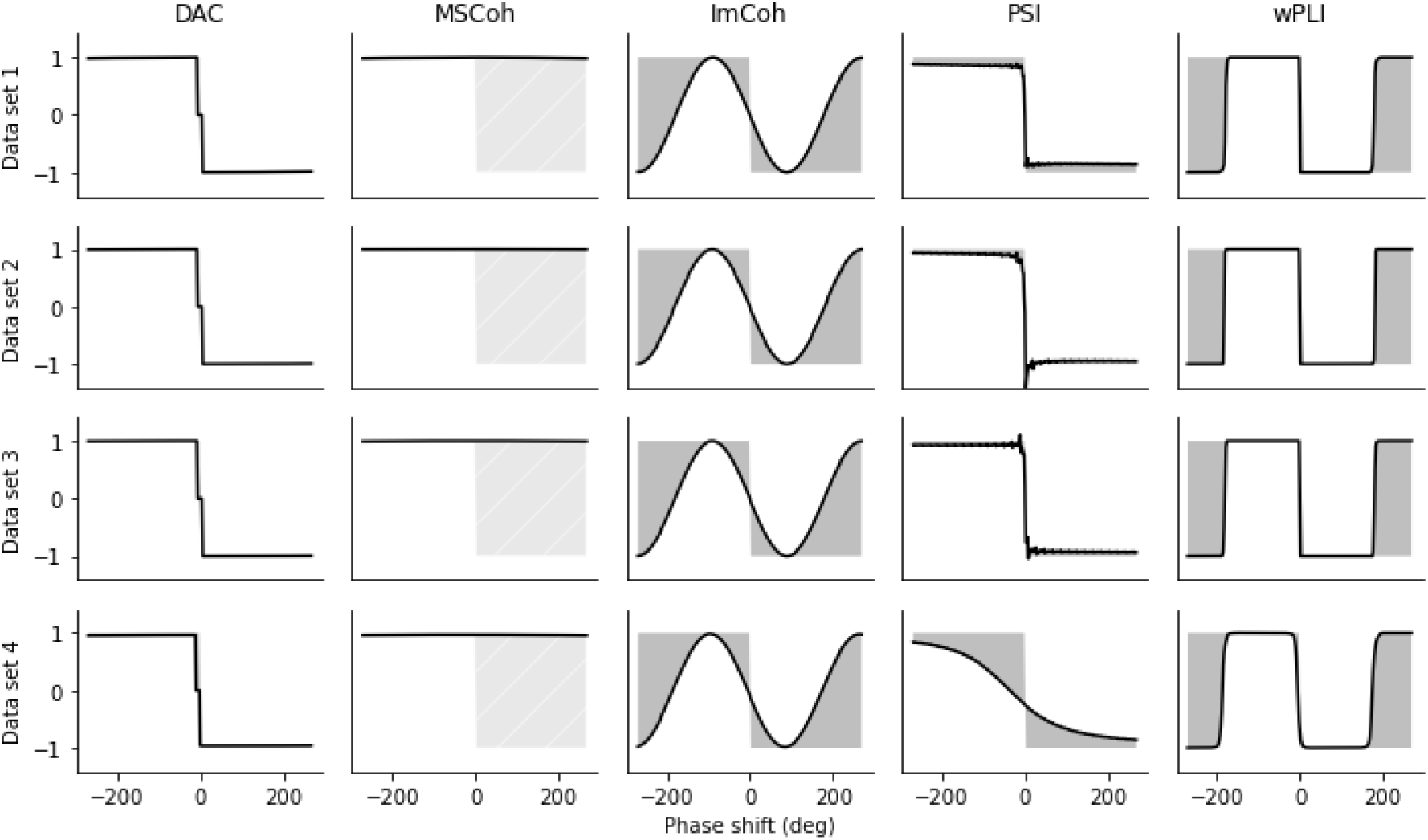
Comparison of connectivity measures. The columns show the performance of directed absolute coherence (DAC), magnitude squared coherence (MSCoh), imaginary coherence (ImCoh), phase slope index (PSI) and weighted phase lag index (wPLI). The areas shaded in grey indicate the difference between the estimated connectivity and the ‘true’ simulated connectivity. Please note that the magnitude squared coherence does not consider the direction of phase shifts and therefore the light grey area, indicating the difference between the estimated magnitude squared coherence and the ‘true’ simulated connectivity, should not be interpreted as the error in the estimated connectivity. For visualization purposes, DAC was modified to return the value zero instead of NaN (Not a Number) in the event of volume conduction.

For data sets 1-3, DAC and magnitude squared coherence correctly estimated the magnitude of the connectivity strength between the original and time-lagged signals, independent of the phase shift. While the modelled maximum connectivity value between the two signals could be detected using imaginary coherency, the absolute maximum value depended on the phase shift between the original and time-lagged signal. This resulted in difficulties in interpretation, since the imaginary coherency at a phase shift of ±90° and ±270° had an absolute value of 1, while the imaginary coherency at, e.g., a phase shift of ±45° and ±225° had an absolute value of only ∼0.72. The normalized PSI provided an accurate estimate of connectivity strength across the entire range of phase shifts, albeit the post-hoc normalization required more data than the other connectivity measures employed. Finally, while wPLI was able to correctly estimate the magnitude of connectivity strength for the majority of the phase shifts, the values proved to be unreliable around phase shifts of 0° and ±180°.

DAC, imaginary coherence, wPLI, and PSI, but not magnitude squared coherence, correctly identified the occurrence of volume conduction at 0° phase shift. However, imaginary coherence and wPLI erroneously identified one case of volume conduction at a phase shift of –180° and +180°, each. Finally, normalized PSI values increased unexpectedly in variability when the phase shift approached 0°. The directionality of the information flow was estimated correctly using DAC and the normalized PSI. Imaginary coherency and wPLI were both able to return information about the directionality for phase shifts between –90° and +90°, but not outside this range.

Both the magnitude and the directionality of the connectivity between the two signals were accurately estimated by DAC for data set 4. A slight decrease in connectivity strength could be observed in comparison to data sets 1-3, as this data set included signals from two different recording sites. A single occurrence of volume conduction was also identified by DAC. Like data sets 1-3, magnitude squared coherence was able to correctly estimate the magnitude of the connectivity strength, but not the presence of volume conduction at 0° phase shift or the directionality of connectivity for phase shifts greater than 0°. Imaginary coherency and wPLI also showed the same issues as observed in data sets 1-3. While successful in identifying the directionality of connectivity, the normalized PSI was unable to correctly determine the magnitude of connectivity strength for data set 4. The absolute maximum values of PSI increased as the lags between the original and time-lagged signal increased, while the modelled level of connectivity remained constant.

## Discussion

The current work presents DAC as a novel measure of effective connectivity. DAC has been designed to be easily interpretable, as it can be explained as the correlation between two signals bound to the absolute interval of -1 and +1. It encapsulates the direction of information flow, is robust against volume conduction, and superior in comparison to several established connectivity measures. Connectivity estimation is a non-trivial task with several pitfalls that require careful consideration if the results are to be interpreted correctly (Bastos and Schoffelen, 2016). One prominent pitfall is the issue of volume conduction. Neuronal signals may propagate more than 1 cm in any direction (Kajikawa and Schroeder, 2011). This issue is further exacerbated for scalp EEG signals which must travel through the skull before being recorded (Nunez and Srinivasan, 2006). When volume conduction occurs, the imaginary part of coherency approaches an absolute value of zero. This effect is used in DAC, imaginary coherence and wPLI to remove effects of volume conduction. However, imaginary coherence and wPLI also approach zero when a phase shift of any integer multiple of 180° occurs. In such conditions, only DAC and PSI were able to detect the phase shift, and demonstrated the highest level of reliability for volume conduction identification amongst the evaluated methods.

Of all the methods under consideration in this study, PSI is the only established measure of effective connectivity. One drawback of the base PSI before normalization is that the estimated values are not bound to a stationary interval. In fact, the bounding interval of base PSI values is proportional to the phase shift between two signals. Therefore, a change in base PSI can be explained by either a phase change, a connectivity change, or a combination of these. This signifies that, although relative changes in connectivity can be quantified, the differences in connectivity between different data sets, or signals acquired over longer time intervals, cannot be so easily compared. In PSI, this issue may be remedied by applying a post-hoc normalization operation such as division by the standard deviation (Nolte et al., 2008), among other potential methods. This requires a larger amount of data, thereby limiting the usability of PSI in clinical settings, e.g., intraoperative recordings, where the amount of data may be limited. Finally, applying post-hoc normalization by means of dividing through the standard deviation approaches a singularity if the phase shift between the investigated signals approaches 0°, as both numerator and denominator approach zero at the same time (see for reference the evaluations of data sets 1-3). In DAC, this issue is remedied by only processing the sign value of PSI, hence discarding any magnitude information.

### Limitations and outlook

Although DAC was able to overcome several disadvantages of magnitude squared coherence, imaginary coherence and PSI by combining the advantages of each individual connectivity method, some limitations remained. Namely, for the volume conduction estimation to be reliable, robust complex connectivity estimates are required. Simultaneously, for the direction identification to be reliable, the frequency bin resolution needs to cover a sufficiently large amount of frequency bins. Both of these limitations are directly inherited from the respective connectivity estimation base methods. Like PSI, DAC is capable of detecting information flow in one direction at a time only. When bidirectional communication occurs between two signals, only the direction with the stronger information flow is identified.

The next steps may include further validation of the proposed method with larger sets of empirical and simulation data from various sources, e.g., collected with surface EEG electrodes. Finally, although the parameters for DAC have been carefully chosen, it is recommended to run the configurator (see supplementary document) to make sure that ones application case aligns with the selected parameters.

## Conclusions

Here, we presented a new method to estimate effective connectivity in neurophysiological data based on the spectral correlation between two signals. DAC is defined between the absolute boundary values of –1 and +1, and is resistant to volume conduction. It borrows a measure of directionality from PSI, while requiring less data to compute. Using local field potential data from patients with Parkinson’s disease, we verified that DAC can provide a more reliable estimate of effective connectivity than other established connectivity measures within the scope of our evaluations. This novel measure may therefore serve as a useful tool for the investigation of brain dynamics in basic and clinical science.

## Acknowledgements

We thank our colleagues at the Institute for Neuromodulation and Neurotechnology for helpful discussions on early versions of the manuscript. This work was supported by the German Federal Ministry of Education and Research (BMBF). We acknowledge support by the Open Access Publishing Fund of the University of Tübingen.

## Code and data availability

The code of this analysis is available at https://github.com/VoodooCode14/dac. The implementation of DAC is available at https://github.com/neurophysiological-analysis/FiNN.

## Credit authorship contribution statement

**Maximilian Scherer**: Conceptualization, Methodology, Software, Writing – original draft, Writing – review & editing.

**Tianlu Wang**: Methodology, Writing – original draft, Writing – review & editing.

**Robert Guggenberger**: Conceptualization, Methodology, Writing – review & editing.

**Luka Milosevic**: Conceptualization, Methodology, Writing – review & editing.

**Alireza Gharabaghi:** Conceptualization, Writing – review & editing, Funding acquisition.

## Supplementary document

A configurator to adjust DAC to a specific recording and investigation combination is available at https://github.com/neurophysiological-analysis/FiNN in the finn_demo/configuration folder. The dac_configuration.py module within this folder can be launched with either sample data from ones analysis or provided demo data. While the module requires the sampling rate, frequency of interest, range around that frequency band, and the desired Fast-Fourier Transform (FFT) resolution (FFT bin width), it may be used to configure the minimal angular threshold (default: 10°) and the volume conductance ratio (default 0.3) to a problem at hand. Although ideal parameters have been carefully selected, optimal results will like require the adjustment of these parameters to one’s investigation. Launching this configurator will evaluate a given parameter set for an artificial phase shift of -270° to +270° around the frequency of interest. This allows one to verify that the provided data & parameter combination yield the expected results.. As data from a single channel is evaluated against a shifted version of itself, volume conductance and the absence thereof can be fully controlled; hence, true volume conduction is expected to only appear around the 0° phase shift.

## Abbreviations

DAC: directional absolute coherence
EEG: electroencephalography
PD: Parkinson’s disease
PLI: phase lag index
PSI: phase slope index
STN: subthalamic nucleus
wPLI: weighted phase lag index

